# Generation time as optimal stopping time

**DOI:** 10.64898/2026.06.30.733937

**Authors:** Tom Froese, Rainer Froese, F. Thomas Bruss

**Affiliations:** Embodied Cognitive Science Unit, Okinawa Institute of Science and Technology Graduate University (OIST), 1919-1 Tancha, Onna-son, Okinawa 904-0495, Japan; Division of Marine Evolutionary Ecology, GEOMAR Helmholtz Centre for Ocean Research Kiel, Kiel, Germany; Department of Mathematics, Faculty of Science, Université Libre de Bruxelles, Brussels, Belgium

**Keywords:** generation time, optimal stopping, 1/*e*-law, life-history theory, reproductive effort, theoretical biology

## Abstract

The timing of peak reproductive effort is a central problem in life-history theory, yet a simple first-principles target for that timing relative to lifespan has been missing. We propose that generation time is an optimal stopping time. Under the 1/*e*-law of best choice, the unique strategy that guarantees the highest minimum probability of selecting a superior option is to wait until a cumulative fraction 1/*e* of opportunities has passed. When reproductive opportunities arrive in an approximately time-homogeneous way over adult life, that threshold is 1/*e* (about 37%) of maximum lifespan. This fraction is confirmed by standard population dynamics for indeterminate growers, combining von Bertalanffy growth with constant mortality. A compilation of published reproductive schedules in plants, animals, and humans illustrates the predicted alignment across taxa and absolute timescales. The 1/*e*-law thus supplies an organizing quantitative account of a core life-history schedule.

## Introduction

Every organism faces a fundamental scheduling problem: when during its lifespan should it invest maximally in reproduction? Reproducing too early sacrifices gains from continued growth, fecundity, or parental expertise, whereas delaying reproduction increases the risk of death before breeding. We propose that this scheduling problem is an instance of optimal stopping, and that generation time is the 1/*e*-law waiting time. A compilation of reproductive schedules from plants to humans illustrates that peak reproductive effort consistently occurs at approximately 1/*e* (∼37%) of species-specific maximum lifespan. This convergence is robust across major phylogenetic groups, absolute lifespan scales, and reproductive modes, indicating a general organizing principle.

The timing of reproductive effort is a central question in life-history theory (Charnov et al. 2007; Roff 1984). Classic work has related age at maturity and peak reproduction to growth, mortality, and fecundity schedules, yet a simple cross-taxa numerical target for the age of peak effort relative to maximum lifespan has been missing. The framework we develop supplies such a target and links it to a known optimum from optimal-stopping theory.

We propose that this regularity reflects an optimal stopping principle formalized by Bruss (1984), a substantial generalization of the classic secretary problem (Gilbert and Mosteller 1966; Lindley 1961). In contrast to the secretary problem, this model requires neither knowledge of the total number of opportunities nor assumptions about when superior options arise. It identifies 1/*e* of the cumulative opportunity distribution as the optimal waiting time and proves that no alternative strategy can guarantee a higher minimum success probability. Applied to biological reproduction, it predicts that organisms should concentrate peak reproductive effort near 1/*e* of lifespan.

If indeed a first-principles framework exists that optimizes the interplay between fecundity, mortality, somatic growth, parental expertise, and probability of success to maximize fitness, then millions of years of natural selection are likely to have implemented it. Our study develops such a framework and illustrates its compatibility with standard population dynamics, compiled life-history records, and the requirements of the generalized optimal stopping theorem (Bruss 1984).

## Materials and methods

### The 1/*e*-law of best choice

Let *N* be a fixed or random positive integer, and let *A_1_*, *A*_2_, …, *A_N_* be a sequence of independent random variables all governed by a common distribution function *F* with density *f* on some interval [0, *t_end_*]. Interpret *N* as a number of uniquely rankable opportunities arriving in [0, *t_end_*], and *t_k_* as the arrival time of rank *k*, with *k* = 1, 2, …, *N*. Suppose that an opportunity can only be selected when it appears (i.e. no recall of preceding opportunities). Let *S*(*t_w_*) be the strategy to wait until time *t_w_*and then to select the first opportunity which ranks better than all preceding ones (if any). Define 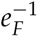 as the time *t* satisfying

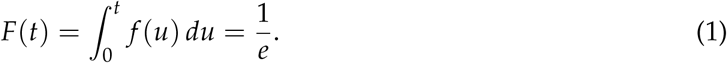

Then the 1/*e*-law of best choice says: The strategy 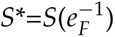 will select rank 1 with probability of at least 1/*e* for all *N* > 0, and *S*\* is the unique strategy which guarantees this result for all *N*. Moreover, 1/*e* is the best possible lower bound for the success probability(Bruss 1984).

The name 1/*e*-law is frequently misused in the literature since non-specialists tend to think it refers to the well-known 1/*e*-rule (37%-rule) for the classical secretary problem (Lindley 1961; Gilbert and Mosteller 1966). Using the waiting period *N*/*e* in the latter, the limiting success probability equals indeed 1/*e* as *N* tends to infinity. Note that this is only a limit result in the special case where *N* is known and where every rank appears equally likely at any arrival time. Other best choice models also need information about *N* (Presman and Sonin 1973). However, for many distributions of *N* the optimal success probability will then fall well below 1/*e* and may even get close to zero. Hence the 1/*e*-law described above came as a true surprise (Samuels 1985).

### Applicability of the 1/*e*-law to biological reproduction

We apply the 1/*e*-law under two operational assumptions developed further in the Discussion. First, in a steady-state population the processes generating objective reproductive opportunities—mates and suitable environmental conditions—are treated as time-homogeneous relative to cohort age, so that their cumulative arrival can be represented, to first approximation, by a uniform distribution over mean maximum lifespan [0, *t_max_*]. Continuous or cyclically gated breeding both yield approximately linear cumulative arrival when averaged over multiple cycles (Dawson et al. 2001; Vasantha 2016). Under that mapping the strategy *S*\* succeeds with probability ≥ 1/*e* at *t_max_*/*e*. Under appreciably non-uniform *F*, the optimum remains at *F*^−1^(1/*e*), which need not equal *t_max_*/*e* (see Fig. 3 and Discussion). Second, actors must be capable of implementing *S*\* through inherited programs, imprinting, learning, or environmental sensing—requirements supported by work in movement, cognitive, and plant ecology and discussed with the empirical pattern below.

### Population dynamics leading up to the 1/*e*-law

Somatic growth is modeled following the Pütter growth equation (Pütter 1920, where the linear dimension *L* is proportional to the cube root of body mass *W* (*L* = *W*^1/3^).

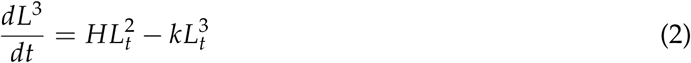

Under approximately isometric growth (W ∝ *L^b^*, with *b* ≈ 3) this formulation can be applied directly to observed body length. von Bertalanffy (1938) reformulated the equation for body mass, yielding the widely used von Bertalanffy growth model

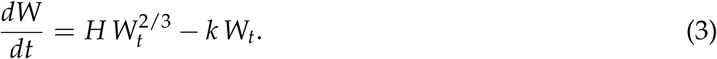

If body mass scales as *W* ∝ *L^b^*, then the relevant exchange surface scales as *L^b^*^−1^. Because *L* ∝ *W*^1/*b*^, the first term in Eq. 3 becomes proportional to *W*^(*b*−1)/*b*^ (Eq. 4).

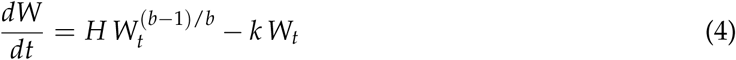

von Bertalanffy (1938) presented the widely used solutions of Eq. 4 for length and weight at age.

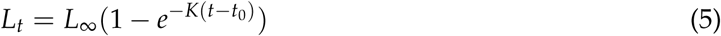

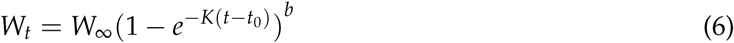

where *L*_∞_ and *W*_∞_ indicate finite asymptotic size, *K* is a growth parameter indicating the exponential decay of the difference to the asymptotic size, *t_0_* is the hypothetical age at zero size, and *b* is the exponent of the weight-at-length equation. In conceptual presentations it is customary to leave out *t_0_* and to assume isometric growth with *b* = 3, as is done below.

For an initial number *N_0_* of cohort members with a constant mortality rate *M*, the number of individuals *N_t_* surviving to a certain age is given by

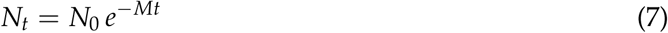

Combining somatic growth with the number of surviving cohort members gives cohort biomass at age as *B_t_* = *N_t_ W_t_*, which reaches a maximum at age *t_Bmax_* (Eq. 8). Since fecundity is about proportional to adult body mass in species with continuous growth, Roff (1984) suggested that the age at maximum cohort biomass *t_Bmax_* would be the optimal age for reproduction.

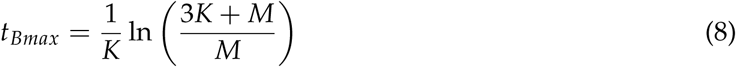

Growth in body weight (Eq. 6 with *b*=3) has an inflection at 8/27 *W*_∞_ corresponding to 2/3 *L*_∞_ at which the production of new tissue is maximum with max *dW/dt* = 4/9 *KW*_∞_ at the corresponding age *t_flex_*.

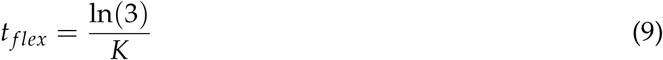

Froese et al. (2016) pointed out that aligning peak reproductive output at *t_Bmax_* with peak tissue production at *t_flex_*reduces the relative energetic cost of reproduction and thus provides a fitness advantage. This alignment is obtained with an *M*/*K* ratio of 3/2. Substituting *K* with 2/3 *M* in Eq. 8 then gives the optimal age for reproduction as *t_opt_*(Jensen 1996).

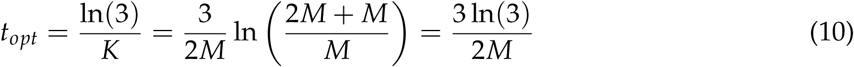

Note that the optimal age for reproduction (Eq. 10 with parameter *M*) also fits observed reproductive schedules of birds and mammals (Table 1, Fig. 3D-F), which cease somatic growth after reaching adult size. If parental competence—essential for successful offspring survival—increases with experience in a saturating manner analogous to the surface-limited scaling (*W*^2/3^) that governs somatic growth, then parameters *K* and *W_t_* would refer to growth of parental competence or expertise and Fig. 1 and the above equations would also apply to birds and mammals.

**Figure 1:**
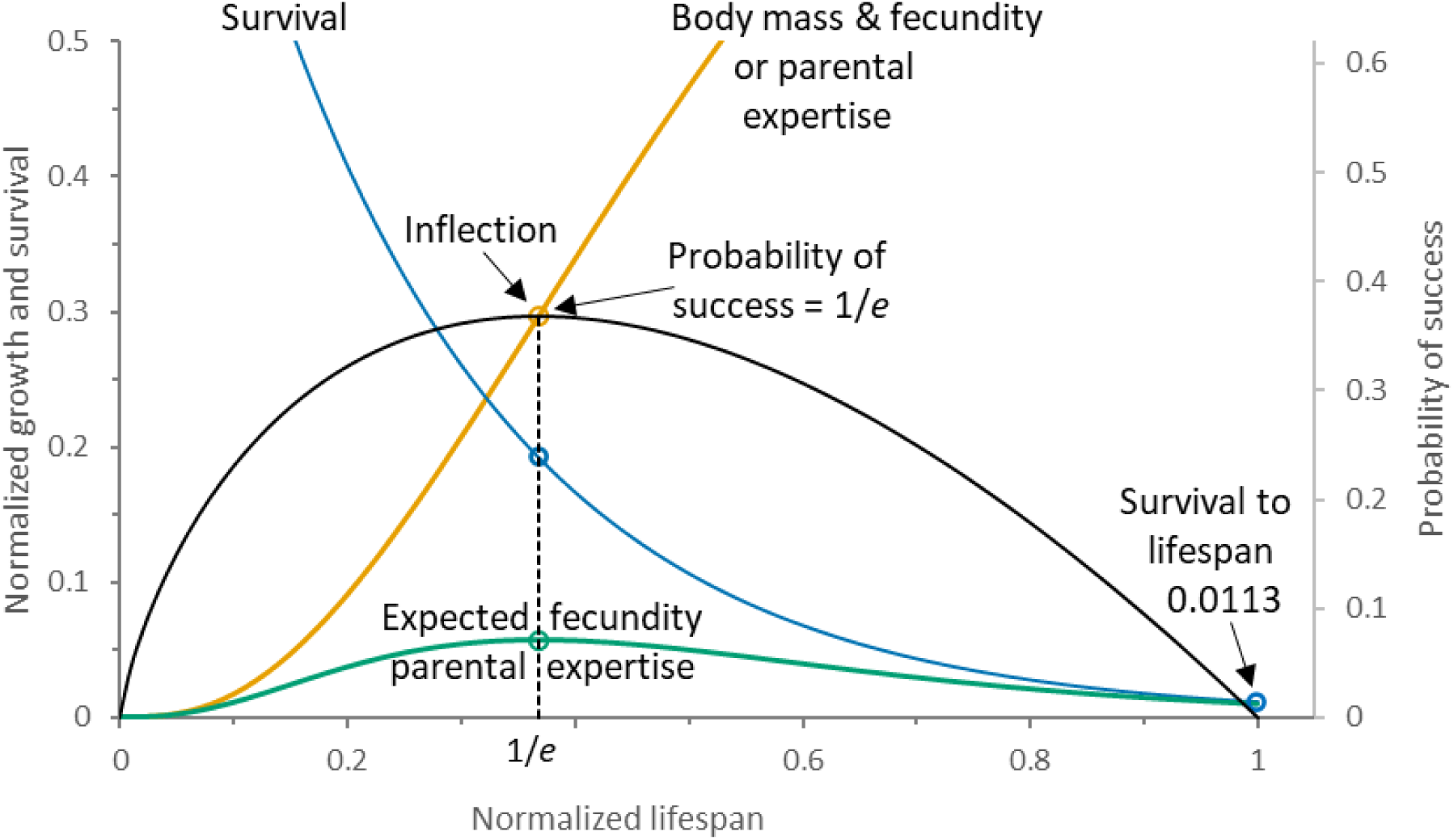
Combining population dynamics with the 1/e-law in a conceptual framework. Survival (blue), somatic growth (orange), and cohort biomass (green, as proxy for expected fecundity or parental expertise) are shown over normalized lifespan. Open dots mark the inflection of the growth curve and the peak of cohort biomass and fecundity. These standard population dynamics are overlaid with the 1/e-law probability of success (black) for a fixed timespan (here: approximate uniform distribution of opportunities mapped on normalized lifespan). The alignment of the age at inflection, and of peak cohort fecundity, with 1/e of lifespan is the empirical regularity analyzed in this study. The height of the success curve is arbitrarily scaled on the second axis such that its peak aligns with the peak of production. For a cohort with such growth and mortality, combining the 1/e-law time frame with the survival curve predicts that about 1% will survive to maximum lifespan.

**Table 1:** Timing of peak reproductive effort relative to lifespan. Exemplary studies of reproductive effort, from viable seed production in trees to births per female per year in humans. Timing of peak reproductive effort overlapped with the predicted ‘optimal window’ of 0.32 – 0.42 of lifespan in all examples.

| Population /<br>Species | N | Indicator | Peak age | Max. age | Peak/Max. | Reference |
| --- | --- | --- | --- | --- | --- | --- |
| European beech<br>tree<br>(*Fagus sylvatica*) | 38 trees | Viable seed<br>production | 148 | 300 – 400 | 0.37 – 0.49 | Bogdziewicz<br>et al. (2023) |
| Wild apple tree<br>(*Malus sieversii*) | 130 trees | Fruit<br>production | 45 – 50 | 130 | 0.35 – 0.38 | Dzhangaliev<br>et al. (2003) |
| Blue mussel<br>(*Mytilus edulis*) | ~ 100<br>mussels | Viable embryo<br>production | 5 | 10 – 14 | 0.36 – 0.5 | Sukhotin and<br>Flyachinskaya<br>(2009)<br>Sukhotin et al.<br>(2007) |
| Spiny lobster<br>(*Panulirus argus*) | NA | Reproductive<br>potential index | 3 - 4 | 10 | 0.3 – 0.4 | Saul (2004)<br>Gnanalingam<br>et al. (2019) |
| Barents Sea cod<br>(*Gadus morhua*) | > 100<br>females | Egg<br>production | 11.7 | 30.5 | 0.38 | Solemdal<br>(1997) |
| European eel<br>(*Anguilla anguilla*) | ~990<br>females | Start of<br>spawning<br>migration | 10 - 14 | 30 – 37 | 0.33 – 0.38 | Simon (2015) |

| Population / Species | N | Indicator | Peak age | Max. age | Peak/Max. | Reference |
| --- | --- | --- | --- | --- | --- | --- |
| Common lizard<br>(*Lacerta vivipara*) | 6 females | Production and survival of offspring | 4 | 11 | 0.36 | Richard et al. (2005) |
| Great tit (*Parus major*) | > 6000 females | Recruit production | 3.5 | 9-10 | 0.35 - 0.39 | Bouwhuis et al. (2009) |
| Red-billed chough<br>(*Pyrrhocorax pyrrhocorax*) | 45 females | Fledgling production | 4 - 6 | 17 - 18 | 0.22 - 0.35 | Reid et al. (2010) |
| African elephants<br>(*Loxodonta africana*) | 834 female elephants | Reproductive prime age of females | 26 | 65 - 74 | 0.35 - 0.4 | Lee et al. (2016) |
| Killer whales<br>(*Orcinus orca*) | > 200 individuals | Viable calf production | 29 - 32 | 80 | 0.36 - 0.4 | Olesiuk et al. (2005) |
| North American red squirrel<br>(*Tamiasciurus hudsonicus*) | 343 females | Recruit production | 2 - 3 | 8 | 0.25 - 0.38 | Descamps et al. (2008) |
| Wild chimpanzees<br>(*Pan troglodytes*) | 17 females | Births/female/year | 20 - 29 | 56 - 61 | 0.33 - 0.52 | Emery Thompson et al. (2007) |
| Aché humans<br>(*Homo sapiens*) | < 324 females | Births/female/year | 30 - 34 | 75 - 80 | 0.38 - 0.45 | Emery Thompson et al. (2007) |
| US humans 2023<br>(*Homo sapiens*) | ~ 3.6 million females | Birth rates by age of mother | 30 - 34 | 82.2 | 0.37 - 0.41 | Osterman et al. (2025)<br>Arias et al. (2025) |

For uniform distributions in the interval [0, 1], the 1/*e*-law of Bruss (1984) of *P*(*x*) = -*x* ln(*x*) gives the probability of success as a function of waiting time *t* as

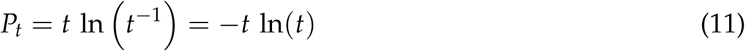

Eq. 11 gives the highest probability of success of *P_t_* = 1/*e* when waiting time is 1/*e* of available time, here assumed equal to lifespan *t_max_*. If natural selection has indeed aligned *t_Bmax_* and *t_flex_* with *t* = 1/*e t_max_*, then with *M*/*K* = 3/2 and *t_max_* = *e t_opt_*, the probability of survival to *t_max_* is obtained from

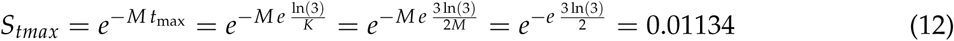

Note that even if the proposal of increase of parental competence with age similar to the somatic growth in weight function is rejected, optimal age for peak reproduction in birds and mammals would still be predicted by the 1/*e*-law to be near *t_opt_* if survival of cohort members to maximum lifespan is about 1%.

### Supporting Evidence

(1) As of March 2026, the von Bertalanffy growth equation has been applied to 13459 populations of 2699 species of fishes (Froese and Pauly 2026) and to 809 marine non-fish species (Palomares and Pauly 2025). Plant growth is commonly described by the Chapman–Richards function, a generalized von Bertalanffy model introduced by Richards (1959). Fitting the von Bertalanffy length equation to single-disk ring-width measurements from 13 trees representing 12 species explained more than 98% of the variance in 11 cases (this study, Table S3), confirming that von Bertalanffy-type growth trajectories can describe long-term growth patterns in trees of up to 760 years. Growth of terrestrial animals is also frequently described by the von Bertalanffy growth equation (Kooijman 2010; Roff 2002; von Bertalanffy 1938).

The 2/3 scaling of the von Bertalanffy growth equation assumes about isometric growth (Eq. 3–4). Froese (2006) examined 3929 published length–weight studies covering 1773 species of fishes and reported a median exponent *b*=3.02 (95% CI 3.011–3.036), with 90% of species having mean *b*-values between 2.7 and 3.4. Comparable analyses of biomass scaling in trees and other vascular plants report exponents typically between about 2.5 and 3.5 (Niklas 1994; Niklas and Enquist 2001). Likewise, in 760 marine species ranging from ctenophores (comb jellies) to marine mammals (seals and whales), 90% of species-specific median *b* estimates fell between 2.4 and 3.3 (this study, using data compiled by (Palomares and Pauly 2025) available in the Supplement). Birds and mammals cease somatic growth after reaching maturity and therefore maintain approximately constant body proportions during adult life.

(2) Figure 2 presents the relative age at first maturity for semelparous and iteroparous fish, invertebrates, birds, mammals and trees. Data on reproductive mode and length at maturation relative to asymptotic length for 1095 populations of 522 species of fishes were obtained from Froese and Pauly (2026). Similar data for 56 populations of 30 invertebrate species were obtained from Palomares and Pauly (2025). The data for relative maturation lengths *L_m_*/*L*_∞_ were converted to relative ages *t_m_*/*t_max_* with

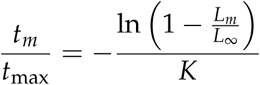

**Figure 2:**
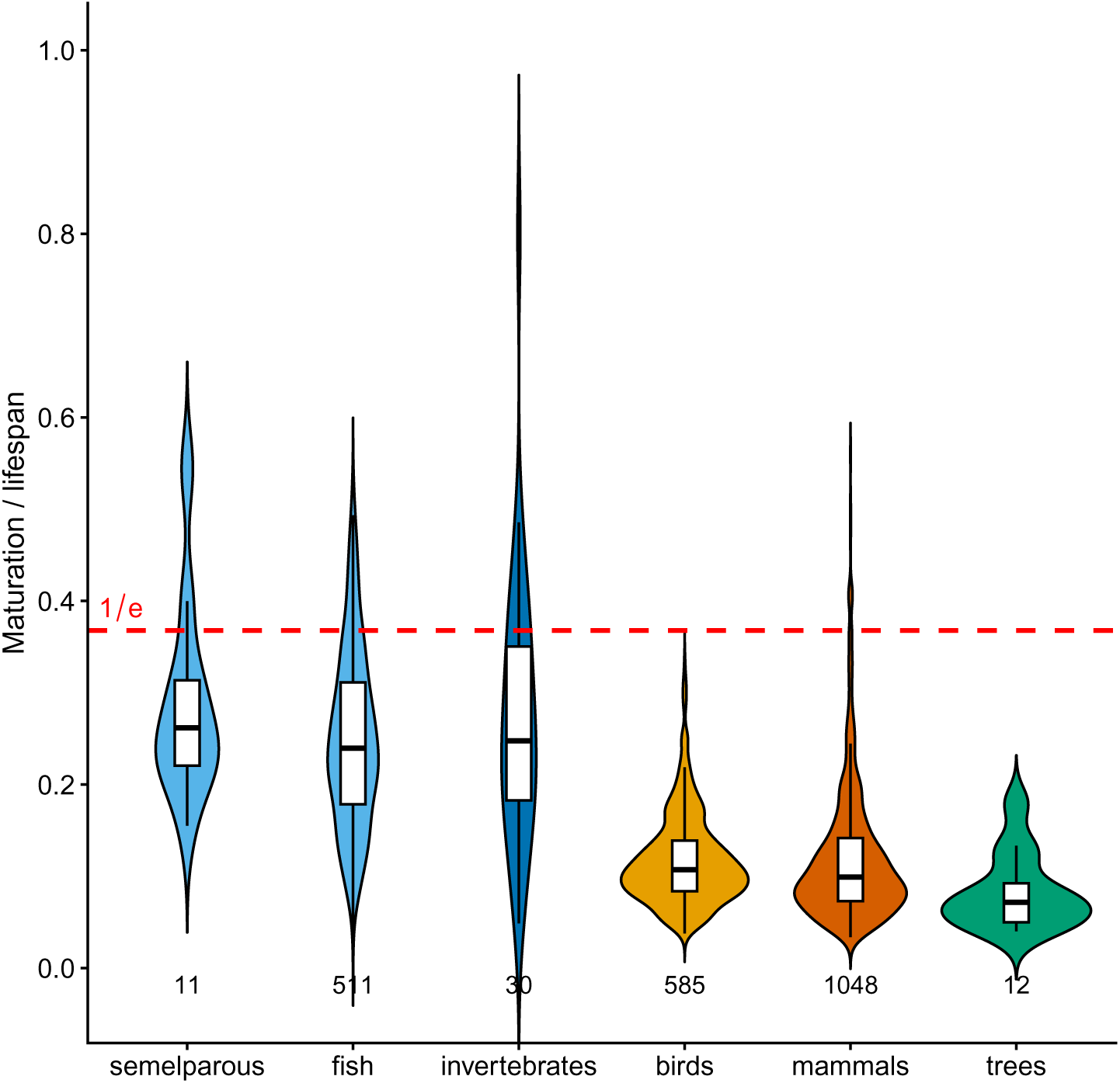
Timing of first maturity relative to lifespan. Violin plots of age at first maturity for 2197 species of semelparous and iteroparous fish, invertebrates, birds, mammals and trees. Birds, mammals and trees use reported ages. Fish use species *K* and wild longevity when longevity is at least the age at 95% of *L*_∞_ (170 species); otherwise the length ratio is converted with *K* set at *t_flex_* = 1/*e* (352 species). Invertebrates use deposited ages (29 species), except one ratio ≥ 1 converted as for the remaining fish. In all groups, maturation tends to occur before 1/e of lifespan (red dashed line). Later maturation is compatible with the 1/e-law but is predicted to be less frequent. Numbers of examined species are indicated.

where *K* was the species-specific growth coefficient when it could be paired with a wild longevity at least the age at 95% of *L*_∞_ (Taylor 1958) (170 fish species). For the remaining 352 fish species, *K* was obtained from Eq. 9 with *t_flex_* = 1/*e*. For 29 invertebrate species, deposited ages were used; one deposited ratio *t_m_*/*t_max_* ≥ 1 was replaced by the same length-ratio conversion. Ratios outside (0, 1) were not plotted. Age at maturation and longevity of 585 species of birds were obtained from (Bird et al. 2020) and for 1048 species of mammals from (Pacifici et al. 2013). Spreadsheets with these data are available in the Supplement. Age at first flowering or first viable seed production and observed longevity were obtained from the literature for 13 trees representing 12 species (Table S3). The mid-ranges of reported maturation age and lifespan were used to calculate *t_m_*/*t_max_*.

(3) The interplay of growth, survival, and the 1/*e*-law (Fig. 1, Eq. 12) predict that about 1% of a cohort will survive to *t_max_*. This prediction is consistent with the empirical data across 202 species, spanning six vertebrate classes and multiple invertebrate taxa (median 1.5%, range 0.9–2.2%) (Dureuil and Froese 2021).

(4) The 1/*e*-law predicts a link between the age at peak reproductive effort and maximum lifespan as *t_opt_* ≈ 1/*e t_max_*. Making use of *t_opt_* ≈ *t_flex_*, we used Eq. 9 to calculate *t_flex_* for 967 species of fishes using species-specific median *K* from Froese and Pauly (2026) (data and R-code available in the Supplement). We then determined the ratio of *t_flex_*to reported maximum ages per species. The median ratio was 0.367, with 50% of the values falling between 0.25 and 0.48. The wide scatter in the data underlines the difficulties in aging and determination of lifespan in wild populations. Table S3 presents a comparison of predicted and observed maximum ages for 12 species of trees, with observed ranges overlapping with predicted ranges in 9 species (75%).

(5) Here we present the data behind the 15 cases of reported timing of peak reproductive effort relative to lifespan, in the sequence of presentation, with common names of species in bold for easier retrieval. Table 1 (Results) summarizes timing of peak reproductive effort relative to lifespan for these cases; source details follow.

1. Bogdziewicz et al. (2023) provide data for viable seed production at size (diameter at breast height in cm, DBH) for 38 **European beech trees** (Fagus sylvatica) sampled from 1980 to 2020, with DBH ranging from about 30 to 170 cm. Data for the three trees with the highest production per reported time period are shown in Table S1. Ages at DBH were derived from Hamilton and Christie (1971) (their Table 71). Maximum age of Beech trees is given as 350 (250 to 400) years (https://ati.woodlandtrust.org.uk/how-to-record/species-guides/beech/) and a possible range of 300 to 400 years was assumed for calculation of relative age.
2. (Dzhangaliev et al. 2003 (p. 206) presents fruit bearing at age for 130 **wild apple trees** of Kazakhstan (*Malus sieversii*). The eldest tree found in the area was aged at 136 years (Dzhangaliev et al. 2003, p. 87) and 130 years lifespan was assumed to calculate relative ages. These trees, described as “representative” rather than randomly selected, may present better-than-average productivity at age, which could explain the surprisingly high productivity at young ages (Fig. 3A).

**Figure 3:**
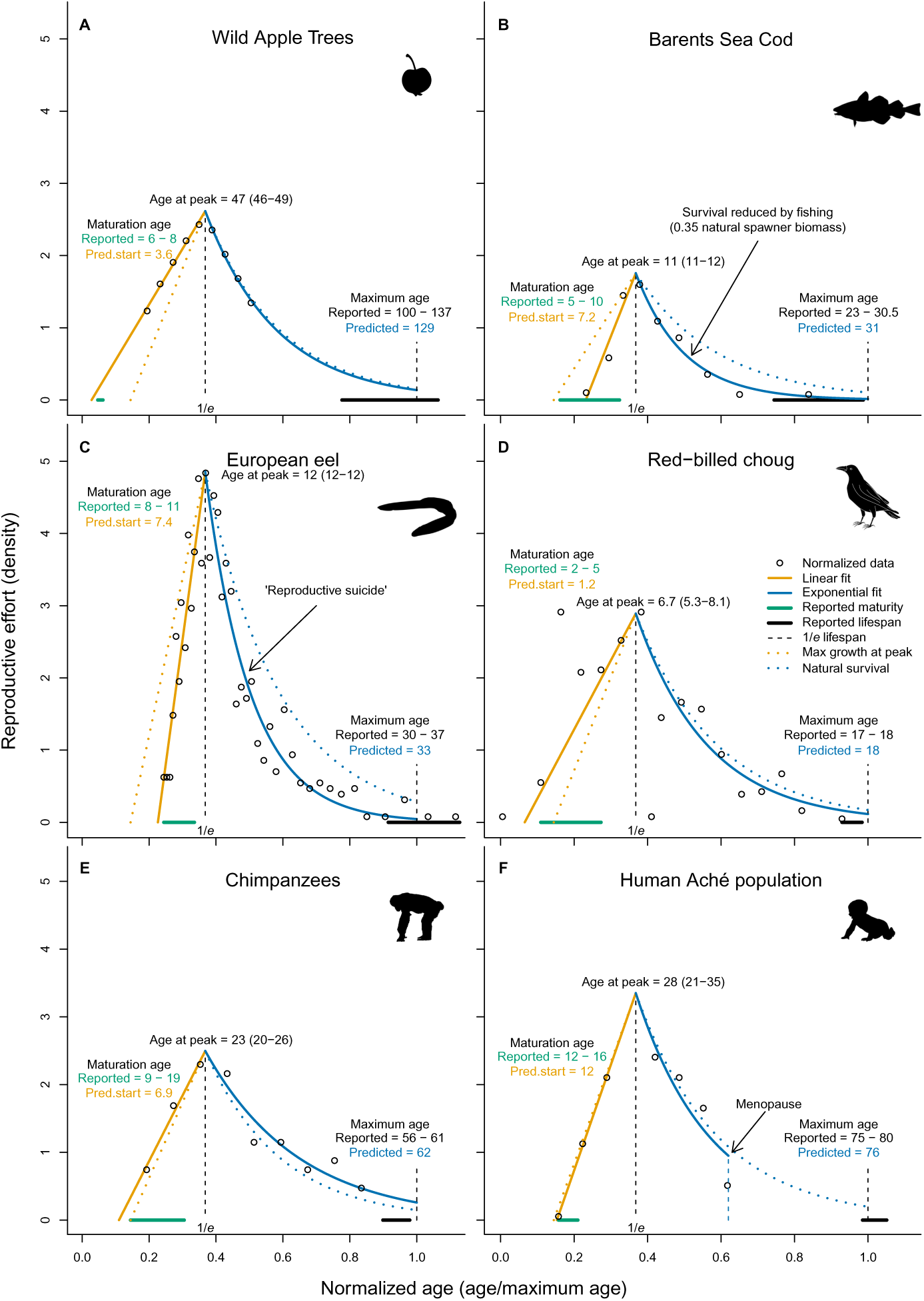
Lifetime reproductive effort of six species interpreted in a 1/e-law framework. Each panel shows normalized reproductive effort (circles) fitted with a piecewise linear-exponential model (solid lines). The dashed vertical line indicates 1/e of maximum age; it is drawn for cross-species visual alignment of peak effort and is not a fitted parameter of the density models. The dotted lines indicate maximum somatic growth rate (left) and population-level survival (right). Reported and predicted maximum ages are indicated. Areas under the curves equal 1, except for Barents Sea cod (0.35), reflecting reduction of spawner biomass by fishing. See Methods for fitting procedure.
3. Sukhotin and Flyachinskaya (2009) present population level and individual viable embryo production for **blue mussel** (*Mytilus edulis*) in the White Sea, with 10 years indicated as maximum age in their Figures. Sukhotin et al. (2007) present maximum individual age of intertidal blue mussels in the White Sea as 14 years.
4. Saul (2004) presents data on population-level fecundity (his ‘Graph 1’) and von Bertalanffy growth curves (his ‘Graph 2’) for female **spiny lobster** (*Panulirus argus*) in the US Virgin Islands. Peak fecundity is reached at 83-88 mm carapace length corresponding to about 3 to 4 years of age. Gnanalingam et al. (2019) give observed maximum age of 10 years.
5. Solemdal (1997) (his Fig. 13) presents data on the egg production of females of **Barents Sea cod** (*Gadus morhua*) based on Norwegian, Russian and Lofoten research cruises from 1986 to 1996. The data for 1992 were standardized and length was converted to age using growth parameters presented in Gislason et al. (2008). The highest age of 30.5 years in the data series was taken as a proxy for *t_max_* and used to normalize age (Fig. 3B). Age at first maturity of 5-10 years was reported by (Godø and Haug 1999). Spawning stock biomass in 1992 was 35% of the highest one in 2013 (ICES 2021), which was taken as a conservative proxy for unexploited biomass and the PDF of reproductive effort was thus standardized to 0.35 instead of 1 as for the other species in Fig. 3.
6. Simon (2015) presents growth parameters for 51 and length frequency data for about 990 female **European eels** captured in the lower Elbe River of Germany in 2011 (Table S1). All females were in the silver eel stage, commencing their migration to the spawning grounds in the Sargasso Sea. Using the indicated mean growth parameters (*L*_∞_=96.1 cm, *K*=0.087, *t*_0_=1.036), a length-age key was constructed. The relative length at maximum tissue production was obtained from *L_flex_*/*L*_∞_ =1 – 1/*b* = 0.684, where *b* is the exponent of the length-weight relationship obtained as 3.16 from 17 studies for females available in Froese and Pauly (2026). That relative length coincided with the mode of length frequencies of the examined silver eel females (Fig. 3C).
7. Richard et al. (2005) present number of surviving offspring of six **common lizard** (*Lacerta vivipara*) females by maternal age, with peak and patterns being identical at the individual and population level. Maximum age of females is given as 11 years.
8. Bouwhuis et al. (2009) (their Fig. 1) present age-specific reproductive performance in female **great tit** (*Parus major*). They report fledgling production by a female of 9 years (their Fig. 2), which was used to assume maximum age of 9-10 years.
9. Reid et al. (2010) (their Fig. 1a) present number of one-year-old offspring by age of 45 females of the **red-billed chough** (*Pyrrhocorax pyrrhocorax*). Maximum age is given as 17 – 18 years (Fig. 3D).
10. Lee et al. (2016) indicate a reproductive “prime age” of 26 years for female **African elephants** (*Loxodonta africana*) in Amboseli National Park. Estimated maximum lifespan is given as 74 years and cohort longevity (95% mortality) as 65 years.
11. Olesiuk et al. (2005) (their Fig. 15) present annual fecundity rate (proportion giving birth to viable calves of either sex) of mature female **killer whales** (*Orcinus orca*) in northern coastal waters of British Columbia as a function of their estimated age during 1997-2004. They also indicate observed maximum age as 80 years.
12. Descamps et al. (2008) (insert in their Fig. 6) report highest number of recruits surviving for one year produced by **North American red squirrel** females of 2 - 3 years of age. Maximum age of females is reported as 8 years.
13. Emery Thompson et al. (2007) (their Table S1) present births per female per year at age of mothers for **wild chimpanzees** (*Pan troglodytes*) of the Kibale (Kanyaeara) population. The youngest to give birth was 14 years old and the oldest 55 years. Age of maturation was deducted from the data as 13-19 years and maximum age of chimpanzees of 56 to 61 years was derived from (Howard-Spink et al. 2025) (Fig. 3E).
14. Births per female per year were obtained for the **human hunter-gatherer Aché** population of Paraguay from Emery Thompson et al. (2007) (their Fig. 1C). Age at menarche was assumed as 12-16 years based on the data and mean maximum lifespan of 75 - 80 years was obtained from Chamberlain (2006) (their Table 2.2) (Fig. 3F).
15. **Human birth rates** by age of mother in the United States for 2023 were obtained from Osterman et al. (2025) (their Table 2). Female life expectancy at age 30 - 34 in 2023 were derived from Arias et al. (2025) (their Table 3).

(6) For the purpose of comparing shapes of lifetime reproductive effort (LRE) distributions across different taxa and life histories, a a segmented linear–exponential regression was fitted to LRE data in a 1/*e* framework. Age data were normalized by division by maximum age and effort was standardized as density, except for Barents Sea cod where the integral under the fit was set to 0.35, according to the natural spawner biomass reduced by fishing (ICES 2021). For comparing the declining fit with population-level survival, the approximate mean hazard rate of the population was obtained from Equation 12 and the corresponding decline in survival is shown as dotted curve in Fig. 3. For comparing the increasing section of the fit with population-level maximum somatic growth *dW/dt* = 4/9 *K W*_∞_, we set *K* = 2/3 *M* and *W*_∞_ = peak density / 0.296. The R-code used for the fit is available in the Supplement. Additional information about the six example species in Fig. 3 is given in Table 1.

## Results

### Standard population dynamics predict an optimal age for reproduction

In organisms with indeterminate growth, fecundity is proportional to adult body mass which follows a von Bertalanffy (1938)-type growth curve (see Methods). For a cohort whose survival is governed by a constant hazard rate, the age at peak cohort biomass and fecundity is a function of the ratio between the hazard rate and the somatic growth rate (Roff 1984). The growth curve has an inflection point where tissue production is maximum and the relative energetic cost of reproduction is lowest, which may provide a fitness advantage for species that align the peaks of tissue- and offspring production (Froese et al. 2016). This proposed optimal age for reproduction *t_opt_* has an unexpected relation with observed maximum lifespan *t_max_* such that *t_max_* ≈ *e t_opt_*, where *e* is Euler’s number (Fig. 1, Table 1).

The 1/*e*-law of Bruss (1984) presents a first-principles explanation to the unexpected *t_opt_*/*t_max_* ≈ 1/*e* ratio. The 1/*e*-law proves that when faced with an unknown number of opportunities of randomly varying quality over a finite timespan, the strategy that maximizes the probability of exploiting the best opportunity is to commit to the first one better than all preceding ones after a well-defined threshold time. This optimal commitment is at 1/*e* of the cumulative opportunity distribution, which coincides with 1/*e* of maximum age under approximately uniform arrival of reproductive opportunities (see Methods). As recently shown also for protozoans (Doan et al. 2026), all organisms sense their environment and act once specific conditions are met. Opportunities for reproduction arrive about uniform during lifespan (see below), missed opportunities cannot be retrieved and future opportunities remain unknown. This fulfils the conditions for the 1/*e*-law and we therefore propose that the empirical 1/*e* ratio found between timing of reproduction and lifespan is not a happenstance but the result of natural selection acting on the optimal timing principle presented by the 1/*e*-law. Below we present relevant evidence.

### Converging strands of evidence

Most scholars define reproductive effort as mass or energy given to reproduction per unit of time, with production of seeds, eggs or offspring often being used as proxy indicator because direct observations of reproductive effort are usually not available (Charnov et al. 2007). Similarly, aging of wild organisms and determination of maximum lifespan are difficult and prone to error. As a result, reliable data on population-level distribution of reproductive effort at age are scarce. Below is a list of summarized direct and indirect evidence in support of the above proposal. Details for each item can be found in the Methods section.

(1) The conceptual framework of population dynamics combined with the 1/*e*-law (Fig. 1) uses a first-principles growth model derived from the geometric scaling of the area through which a volume of body tissue exchanges mass and energy with its environment (Pütter 1920; von Bertalanffy 1938). The model assumes about isometric growth of body proportions (mass ∝ length*^b^*, with 2.5 < *b* < 3.5), which is indeed predominant across taxa. The growth model has been successfully fitted to thousands of species from plants to mammals.

(2) A necessary condition for the predicted optimal age of peak reproductive effort *t_opt_* ≈ 1/*e t_max_* is that, on average, maturation occurs mostly at or before *t_opt_*. Figure 2 presents the relative age at first maturity for 2197 species ranging from trees to mammals. In 96.0% of the species examined, maturation occurred at or before 1/*e* of lifespan (Fig. 2).

(3) Survival, growth and cohort biomass curves are asymptotic (Fig. 1), permitting prediction of the optimal reproductive age, *t_opt_*, but not of lifespan *t_max_*. However, if natural selection has aligned peak reproductive effort with the 1/*e*-law optimum, then a survival probability to maximum age of 1.1% is predicted, which is compatible with published data (Dureuil and Froese 2021).

(4) A testable key prediction of the 1/*e*-law is that peak reproductive effort occurs at about 1/*e* of maximum age. Taylor (1958) suggested that in species with indeterminate growth the age at 95% of asymptotic length was a good approximation of maximum age. Comparing that with the age at inflection of the growth curve gives a previously unnoticed ratio of about 1/*e* (∼37%), which triggered the research leading to this study. We compared median age at inflection to reported maximum age for 967 species of fishes and found a median ratio of 0.37, with 50% of the observations between 0.25 and 0.48.

(5) A literature search across major phylogenetic groups for direct observations of reproductive effort by parental age found 15 exemplary studies from plants to humans, with maximum ages from 8 to about 400 years (Table 1). Reported peak reproductive ages overlapped with an ‘optimal timing’ window defined as 0.32 < 1/*e* < 0.42 of lifespan, where the 1/*e*-law success probability is ≥ 0.99/*e*.

(6) Charnov et al. (2007) proposed that lifetime reproductive effort (LRE) relative to adult body mass is approximately constant across organisms. We fitted segmented linear–exponential density distributions to LRE data for six species spanning trees, fish, birds, and primates (Fig. 3). The distributions share a common structure: after maturation reproductive effort rises steeply to a peak near 1/*e* of lifespan, with a slope close to the maximum growth rate at inflection. It then declines exponentially, tracking population-level survival. The shapes vary predictably: the semelparous eel (Fig. 3C) shows the highest density and narrowest width, reflecting total conversion of somatic resources into a single reproductive event followed by death (‘reproductive suicide’), while the long-lived apple tree (Fig. 3A) shows the widest distribution. The strongly exploited Barents Sea cod (Fig. 3B) is an expected (regrettable) outlier, with fishing mortality steepening the decline in effort below population-level survival (ICES 2021). Despite their fast somatic growth reaching terminal size near early maturation (Fig. 2), birds and mammals (Fig. 3D-F) adhere to the same lifetime-distribution of reproductive effort around 1/*e* of lifespan as the other species. Note that the reproductive phase is always shorter than lifespan, especially in semelparous species (Fig. 3C) and species with menopause (Fig. 3F). But the placement of the reproductive phase within lifespan is such that its peak aligns again with the predicted optimum at 1/*e* of lifespan. The distribution of reproductive effort within the reproductive phase appears to be also governed by the 1/*e*-law with cumulative density before the peak varying around the predicted 0.37 (0.32 – 0.44; Fig. 3) of total effort.

### Semelparous species, iteroparous species, birds and mammals, and humans

Semelparous species with good estimation of maximum lifespan, such as available for the European eel (*Anguilla anguilla*), map easiest with the proposed concept of evaluating opportunities for reproduction and taking the best so far after about 1/*e* lifespan of waiting and observing (Fig. 3C). Iteroparous species spread reproductive events across their reproductive phase, however, with a clear high peak or narrow plateau of population-level reproductive effort. This peak or plateau could be anywhere, early or late in life, but instead clusters around 1/*e* of lifespan (Table 1, Fig. 3), similar to the semelparous case. This may surprise, but should no longer, because Roff (1984) showed that the optimal age of reproduction was the same for semelparous and iteroparous species. More generally, the shape of the population-level lifetime reproductive effort distribution is unimodal and the optimal timing of the mode is a unique lifetime decision independent of the width of the distribution.

Most birds and mammals approach their maximum size at about 1/10th of their lifespan and mature shortly after (Fig. 2). Their clutch or litter size is decoupled from body size and varies around a species-specific constant. However, their fledgling or surviving offspring production is not immediately declining with population-level survival as suggested by constant fecundity, but instead clusters also around 1/*e* of lifespan (Table 1, Fig. 3D-F). For example, our largest examined data set analysed age-at-giving-birth of 3.6 million human mothers and—despite medical and societal confounds—peaked close to the predicted 1/*e* ratio (Table 1).

## Discussion

Across plants, invertebrates, fishes, birds, mammals, and humans, peak reproductive effort clusters near 1/*e* of species-specific maximum lifespan. That alignment is the empirical illustration of the theoretical claim: generation time is an optimal stopping time. Two independent lines of argument land on the same fraction. Standard population dynamics for indeterminate growers, with cohort biomass peaking when mortality and somatic growth are aligned (*M*/*K* ≈ 3/2), predict *t*_opt_/*t*_max_ ≈ 1/*e*. Independently, the 1/*e*-law of best choice (Bruss 1984) proves that, when an unknown number of rankable opportunities arrives over a finite horizon, the strategy that guarantees the highest minimum probability of selecting the best is to wait until a cumulative fraction 1/*e* of opportunities has passed and then take the first candidate better than all previous ones. Under approximately uniform arrival of reproductive opportunities over lifespan—an inference supported by the biology of continuous or cyclically gated breeding and by the data themselves—that threshold falls at *t*_max_/*e*. The theorem is not a metaphor for selection; it is a proven optimum under well-specified conditions. That biology so consistently places peak effort at the same fraction is strong evidence that natural selection has implemented a schedule that realizes this optimum (Bruss 1984).

The mapping from abstract opportunities to biological reproduction rests on time-homogeneous arrival and on the capacity to implement the stopping strategy. Continuous or seasonally gated breeding supplies approximately equal numbers of opportunities per unit time when averaged over cycles, so cumulative arrival is near-linear over lifespan (Dawson et al. 2001; Vasantha 2016). The empirical peak of effort near *t*_max_/*e* itself supports that uniformity: under strongly non-uniform *F*, the optimum would still be at *F*^−1^(1/*e*) but would not necessarily coincide with *t*_max_/*e*, as illustrated by within-phase peaks around 1/*e* of cumulative density in Fig. 3. Actors need not “know” the theorem: animals use inherited orientation programs, early-life imprinting, and learning from environmental and social cues to time migration and breeding (Danchin et al. 2004; Kashetsky et al. 2021), and plants use environmental sensing and memory (e.g. vernalization and masting) (Bogdziewicz et al. 2023). Those mechanisms, together with roughly uniform opportunity arrival, satisfy the conditions of optimal-stopping theory for lifespan timing of reproduction.

The integral condition *F*(*t*) = 1/*e* is distribution-general: it identifies the unique strategy that guarantees success probability at least 1/*e* for every *N >* 0, and thus underpins the predominance of best and good selection actions under the theorem’s hypotheses (Bruss 1984). Stronger procedures, notably the odds-algorithm, can raise success probability further when *N* is known or estimable (Bruss 2000). Biological reproduction typically lacks that information, so the 1/*e*-law is the appropriate bound for the cross-taxon schedule reported here. Where *N* can be estimated in particular systems, those stronger results become available for applied work and exploit more of the same mathematical generality.

### Why was this pattern not identified earlier?

Reliable joint observations of age-specific reproductive effort and maximum lifespan are scarce: aging wild individuals is difficult, and direct measures of reproductive effort by age are rarer still (Charnov et al. 2007). Much life-history work has focused on age at maturity or on generation time defined in other ways, rather than on the mode of the lifetime effort distribution relative to *t*_max_. The present study is not a small cherry-picked gallery. The maturity screen spans **2,197 species** from trees to mammals (96.0% mature at or before 1/*e* of lifespan; Fig. 2). The growth-inflection ratio is assessed in **967 species of fishes** (median *t*_flex_/*t*_max_ = 0.367). Exemplary effort-by-age series (Table 1; Fig. 3) cover orders-of-magnitude differences in absolute lifespan. The pattern survives that breadth.

### When do species depart from *t*_opt_?

Table 1 and Fig. 3 also show species whose peak effort sits somewhat above or below 1/*e*. Such departures are expected when the model’s conditions are strained. Heavy exploitation can truncate the older age classes and shift or compress the effort distribution, as in Barents Sea cod under fishing mortality (ICES 2021). Menopause and other post-reproductive life (e.g. humans) shorten the reproductive window relative to total lifespan while the peak within that window can still align with 1/*e* of maximum age. Semelparous species concentrate effort in a narrow burst; the mode still tracks the predicted optimum even when the distribution is narrow. Measurement error in *t*_max_ and in aging widens scatter—visible in the fish inflection ratios (IQR 0.25–0.48)—without erasing the central tendency at 0.37. In biological terms, we expect the 1/*e* schedule when reproductive opportunities are available in a roughly time-homogeneous way over adult life, missed chances cannot be revisited, and fitness benefits of waiting (growth, fecundity, or parental skill) saturate while mortality continues. Where opportunity arrival is strongly front- or back-loaded, the general law still places the optimum at *F*^−1^(1/*e*), which need not equal *t*_max_/*e*; the within-phase peaks in Fig. 3 illustrate that case.

## Conclusions

The convergence of growth physiology, peak reproductive effort, and optimal stopping on the same 1/*e* fraction of lifespan explains a previously unrecognized quantitative regularity across taxa. It also yields a first-principles prediction for maximum lifespan with an associated survival-to-maximum-age of about 1.1% that matches empirical estimates across vertebrate and invertebrate taxa (Dureuil and Froese 2021). The 1/*e*-law framework adds a first-principles target for natural selection and a practical means to infer otherwise inaccessible life-history parameters from minimal information, with direct applications to the conservation and management of exploited or threatened populations.

## Acknowledgments

We thank Mark J. Costello for his comments on an earlier version of the manuscript.

## Author Contributions

RF noticed the 1/*e* ratio between peak reproductive effort and lifespan, initiated this study and wrote the first draft text. TF proposed the optimal waiting time theorem as the explanation of the observed ratio, finalized the write-up and submitted the paper as corresponding author. RF and TF invited FTB to discuss in detail the many subtleties and implications of the best-choice objective and the 1/e-law.

## Data and Code Availability

Data and R scripts supporting the findings are available in the online repository at https://oceanrep.geomar.de/id/eprint/63525/ (GEOMAR OceanRep; CC-BY). Additional tables appear in the Supplemental Information.

