## Supplementary material for "Generation time as optimal stopping time": Online Supplement

Tom Froese[1,*]

Rainer Froese[2]

F. Thomas Bruss[3]

### Extended Data

**Table S1:** Table S1. Diameter at breast height (DBH) of individual European Beech trees with the highest production of viable seeds (Ranks 1-3) during the indicated time intervals. The age corresponding to DBH is given together with the ratios of age to 300 or 400 years maximum lifespan.

| Years | DBH 1 | DBH 2 | DBH 3 | Mean DBH | Age | Age/300 | Age/400 |
|---|---|---|---|---|---|---|---|
| 1980-1990 | 72.9 | 74.0 | 52.1 | 66.3 | 117 | 0.39 | 0.29 |
| 1985-1995 | 66.0 | 62.4 | 95.0 | 74.5 | 133 | 0.44 | 0.33 |
| 1990-2000 | 69.0 | 98.6 | 128.2 | 98.6 | 181 | 0.60 | 0.45 |
| 1995-2005 | 81.4 | 72.0 | 68.0 | 73.8 | 139 | 0.46 | 0.35 |
| 2000-2010 | 58.2 | 72.5 | 105.1 | 78.6 | 142 | 0.47 | 0.35 |
| 2005-2015 | 85.0 | 108.0 | 68.0 | 87.0 | 158 | 0.53 | 0.40 |
| 2010-2020 | 87.9 | 86.4 | 110.0 | 94.8 | 173 | 0.58 | 0.43 |
| Mean | | | | 81.9 | 148 | 0.49 | 0.37 |

**Table S2:** Table S2. Length, age and frequency of silver eel (Anguilla anguilla) females caught in the lower Elbe River at the start of their spawning migration. Mean growth parameters were used to transform length to age. The age at 95% of asymptotic length (33.9 years for $L_\infty$ = 96.1 cm, K = 0.087, $t_0$ = 1.04) was used as proxy for maximum age.

| Length (cm) | $L/L_\infty$ | Age (years) | Age / max age | Frequency |
|---|---|---|---|---|
| 52.5 | 0.55 | 8.0 | 0.24 | 8 |
| 53.6 | 0.56 | 8.3 | 0.25 | 8 |
| 54.5 | 0.57 | 8.6 | 0.26 | 8 |
| 55.5 | 0.58 | 8.9 | 0.27 | 19 |
| 56.5 | 0.59 | 9.2 | 0.27 | 33 |
| 57.6 | 0.60 | 9.5 | 0.28 | 25 |
| 58.4 | 0.61 | 9.7 | 0.29 | 39 |
| 59.5 | 0.62 | 10.1 | 0.30 | 31 |
| 60.5 | 0.63 | 10.4 | 0.31 | 51 |
| 61.5 | 0.64 | 10.7 | 0.32 | 38 |
| 62.4 | 0.65 | 11.0 | 0.33 | 48 |
| 63.5 | 0.66 | 11.4 | 0.34 | 61 |
| 64.5 | 0.67 | 11.7 | 0.35 | 46 |
| 65.5 | 0.68 | 12.1 | 0.36 | 62 |
| 66.4 | 0.69 | 12.5 | 0.37 | 47 |
| 67.4 | 0.70 | 12.9 | 0.39 | 58 |
| 68.5 | 0.71 | 13.3 | 0.40 | 55 |
| 69.5 | 0.72 | 13.7 | 0.41 | 40 |
| 70.3 | 0.73 | 14.1 | 0.42 | 46 |
| 71.4 | 0.74 | 14.6 | 0.44 | 41 |

| Length (cm) | $L/L_\infty$ | Age (years) | Age / max age | Frequency |
|---|---|---|---|---|
| 72.5 | 0.75 | 15.1 | 0.45 | 21 |
| 73.4 | 0.76 | 15.6 | 0.47 | 24 |
| 74.5 | 0.78 | 16.1 | 0.48 | 22 |
| 75.4 | 0.78 | 16.6 | 0.50 | 25 |
| 76.4 | 0.80 | 17.2 | 0.51 | 14 |
| 77.5 | 0.81 | 17.8 | 0.53 | 11 |
| 78.4 | 0.82 | 18.4 | 0.55 | 17 |
| 79.3 | 0.83 | 19.0 | 0.57 | 9 |
| 80.4 | 0.84 | 19.8 | 0.59 | 20 |
| 81.4 | 0.85 | 20.6 | 0.62 | 12 |
| 82.4 | 0.86 | 21.4 | 0.64 | 7 |
| 83.5 | 0.87 | 22.3 | 0.67 | 6 |
| 84.5 | 0.88 | 23.3 | 0.70 | 7 |
| 85.5 | 0.89 | 24.3 | 0.73 | 6 |
| 86.4 | 0.90 | 25.4 | 0.76 | 5 |
| 87.5 | 0.91 | 26.7 | 0.80 | 6 |
| 88.4 | 0.92 | 27.9 | 0.84 | 1 |
| 89.4 | 0.93 | 29.7 | 0.89 | 1 |
| 90.5 | 0.94 | 31.6 | 0.95 | 4 |
| 91.5 | 0.95 | 33.9 | 1.01 | 1 |
| 92.5 | 0.96 | 36.6 | 1.10 | 1 |
| 93.7 | 0.97 | 41.2 | 1.23 | 1 |
| 95.4 | 0.99 | 55.8 | 1.67 | 1 |

**Table S3:** Table S3. Von Bertalanffy growth in length was fitted to the longest time series of individual tree ring widths for all species in GenTree (`https://www.gentree-h2020.eu/`) and selected additional species in ITRDB (`https://www.ncei.noaa.gov/products/paleoclimatology/tree-ring`), with indication of the tree-ID, ring count (~ age), coefficient of determination $r^2$, growth parameter K, and the predicted maximum age. Values in parentheses are 95% confidence limits for K and application of the lower and upper bounds of the 'optimal timing' window around 1/e, viz. 0.32 – 0.42. Published ranges for lifespan and record age are indicated, with sources. Note that ring counts (bold) of two trees set new age records for the species. Predicted ranges that overlap with published ranges are marked bold.

| Species | ITRDB/ Gen-Tree | Ring count | $r^2$ | $K$ | Predicted max age | Maturation age | Lifespan | Record age | Sources |
|---|---|---|---|---|---|---|---|---|---|
| *Betula pendula* | FRBP2113 | 88 | 0.983 | 0.0097 (0.007 - 0.012) | 284 (247 - 324) | 10 - 20 | 100 - 150 | ~235 | 1, 2, 13 |
| *Cedrela odorata* | MEXI049 - ARN05C | **251** | 0.997 | 0.0092 (0.009 - 0.0095) | 330 (287 - 376) | 10 - 15 | 84 – 250 | ~255 | 3, 4, 14 |
| *Fagus sylvatica* | FRFS0617 | 285 | 0.986 | 0.0049 (0.0046 - 0.0053) | 643 (**559 - 733**) | 30 - 50 | 250 - 350 | ~**625** | 1, 5, 15, 17 |
| *Picea abies* | DEPA1011 | 214 | 0.992 | 0.011 (0.01 - 0.011) | **288 (250 - 328)** | 20 - 40 | **250 - 400** | ~468 | 1, 6, 13, 18 |

| Species | ITRDB/ Gen-Tree | Ring count | $r^2$ | $K$ | Predicted max age | Maturation age | Lifespan | Record age | Sources |
|---|---|---|---|---|---|---|---|---|---|
| *Picea sitchensis* | AK121 - SIT121A | 593 | 0.989 | 0.0039 (0.0038 - 0.0041) | **665 (579 - 758)** | 20 - 40 | **500 - 700** | **>700** | 1, 6, 15 |
| *Pinus pinaster* | FRPP1408 | 120 | 0.991 | 0.023 (0.021 - 0.024) | 130 (113 - 149) | 10 - 20 | 150 - 300 | ~409 | 1, 9, 19 |
| *Pinus sylvestris* | GBPS1304 | 325 | 0.997 | 0.0052 (0.0051 - 0.0054) | **581 (506 - 663)** | 15 - 30 | **300 - 600** | ~757 | 1, 6, 13 |
| *Populus nigra* | DEPO1009 | 109 | 0.961 | 0.025 (0.024 - 0.026) | **106 (92 - 121)** | 10 - 15 | **100 - 200** | ~400 | 1, 10, 19 |
| *Pseudotsuga menziesii* | CO021 - 642143 | 760 | 0.992 | 0.0034 (0.0033 - 0.0035) | **879 (765 - 1002)** | 20 - 40 | **500 - 1000** | ~1333 | 1, 6, 13 |
| *Pseudotsuga menziesii* | OR123 - HAN01A | 169 | 0.995 | 0.013 (0.013 - 0.014) | 234 (204 - 267) | 20 - 40 | 500 - 1000 | ~1333 | 1, 6, 13 |
| *Quercus petraea* | ITQP0912 | 319 | 0.998 | 0.0043 (0.0041 - 0.0044) | **673 (585 -767)** | 40 - 60 | **500 - 800** | ~1000 | 1, 11, 16 |

| Species | ITRDB/ Gen-Tree | Ring count | $r^2$ | $K$ | Predicted max age | Maturation age | Lifespan | Record age | Sources |
|---|---|---|---|---|---|---|---|---|---|
| *Quercus robur* | GERM011 - 371041 | 318 | 0.993 | 0.0032 (0.003 - 0.0034) | **942 (819 - 1073)** | 40 - 60 | **500 - 1000** | ~1200 | 1, 11, 16 |
| *Tectona grandis* | INDO007 - SARA02C | **307** | 0.932 | 0.0075 (0.0072 - 0.0078) | **345 (300 - 394)** | 10 - 20 | 200 - 400 | ~500 | 3, 12, 14 |

**Table S4:** Table S4. References used as source in Table S3.

| **References** |
|---|


[1] Burns, R. M. & Honkala, B. H. (Eds.). (1990). Silvics of North America: Volume 1. Conifers; Volume 2. Hardwoods (Agriculture Handbook 654). U.S. Department of Agriculture, Forest Service.

[2] Atkinson, M. D. (1992). Betula pendula Roth (B. verrucosa Ehrh.). Journal of Ecology, 80(4), 837–870.

[3] Orwa, C., Mutua, A., Kindt, R., Jamnadass, R., & Anthony, S. (2009). Agroforestree Database: A Tree Reference and Selection Guide. World Agroforestry Centre (ICRAF). http://www.worldagroforestry.org/treedb2/

[4] Lamb, F. B. (1968). Mahogany of Tropical America: Its Ecology and Management. Berkeley: University of California Press.

[5] Ellenberg, H., & Leuschner, C. (2010). Vegetation of Central Europe (5th ed.). Cambridge: Cambridge University Press.

[6] European Forest Institute / EUFORGEN species accounts.
